# BMP-signalling inhibition in *Drosophila* secondary cells remodels the seminal proteome, and self and rival ejaculate functions

**DOI:** 10.1101/741587

**Authors:** Ben R. Hopkins, Irem Sepil, Sarah Bonham, Thomas Miller, Philip D. Charles, Roman Fischer, Benedikt M. Kessler, Clive Wilson, Stuart Wigby

## Abstract

Seminal fluid proteins (SFPs) exert potent effects on male and female fitness. Rapidly evolving and molecularly diverse, they derive from multiple male secretory cells and tissues. In *Drosophila melanogaster*, most SFPs are produced in the accessory glands, which are composed of ∼1000 fertility-enhancing ‘main cells’ and ∼40, more functionally cryptic, ‘secondary cells’. Inhibition of BMP-signalling in secondary cells suppresses secretion, leading to a unique uncoupling of normal female post-mating responses to the ejaculate: refractoriness stimulation is impaired, but offspring production is not. Secondary cell secretions might therefore make a highly specific contribution to the seminal proteome and ejaculate function; alternatively, they might regulate more global – but hitherto-undiscovered – SFP functions and proteome composition. Here, we present data that supports the latter model. We show that in addition to previously reported phenotypes, secondary cell-specific BMP-signalling inhibition compromises sperm storage and increases female sperm use efficiency. It also impacts second male sperm, tending to slow entry into storage and delay ejection. First male paternity is enhanced, which suggests a novel constraint on ejaculate evolution whereby high female refractoriness and sperm competitiveness are mutually exclusive. Using quantitative proteomics, we reveal a mix of specific and widespread changes to the seminal proteome that surprisingly encompass alterations to main cell-derived proteins, indicating important cross-talk between classes of SFP-secreting cells. Our results demonstrate that ejaculate composition and function emerge from the integrated action of multiple secretory cell-types suggesting that modification to the cellular make-up of seminal fluid-producing tissues is an important factor in ejaculate evolution.

## INTRODUCTION

Ejaculates are compositionally rich. In addition to sperm, males transfer a cocktail of proteins (seminal fluid proteins, ‘SFPs’), lipids, salts, vesicles, and nucleic acids, which together constitute the seminal fluid (1–3). The phenotypic effects of seminal fluid in females are broad, particularly in invertebrates. In various species these effects include increased aggression, reduced sexual receptivity, shifts in dietary preference, conformational changes in the reproductive tract, immuno-modulation, and stimulation of offspring production (reviewed in 4–6). A number of SFPs have been further implicated in sperm competition, the battle between sperm from different males for fertilisations (7–10). Consequently, seminal fluid represents a critical mediator of male reproductive success (11, 12).

While sperm are always produced in testes, seminal fluid generally comprises products drawn from a number of reproductive tissues (13). These tissues vary considerably in number, cellular make-up, and developmental identity between species, with lineages showing evolutionary patterns of loss, modification, and acquisition (4, 13–15). Why male reproductive systems incorporate this diversity is unclear. It has been suggested that by sequestering SFPs in different cells or glands males are afforded control over their release, and consequently, spatiotemporal control over their interactions with sperm, the female reproductive tract, or with other SFPs (16). Additionally, functional diversification of tissues and cell-types may be required to build specialised parts of the ejaculate, such as mating plugs (17). In either case, activities may be carried out independently between cell-types and tissues or there may be cross-talk between them that coordinates global seminal fluid composition. Such cross-talk may be required to drive the sophisticated strategic changes in ejaculate composition observed in relation to sperm competition threat (18). Fundamentally, to understand how ejaculates evolve it is essential that we understand the drivers of diversity in the elements within the male reproductive system, as well as the functional connectivity between them.

The male reproductive system of *Drosophila melanogaster* consists of testes that produce sperm, and three secretory tissues that contribute to the seminal fluid: the paired accessory glands, ejaculatory duct, and ejaculatory bulb (4)(Fig. 1A). The majority of the ∼200 SFPs known to be transferred to females are produced and stored in the accessory glands (19). Each of the two lobes of the glands is composed of two distinct cell-types (20). The majority are the ∼1000 small, binucleate ‘main cells’ (20), which are thought to produce most of the gland’s secretion (21). Accordingly, these cells have been shown to be the sole production site for several highly-abundant and functionally-important SFPs, including sex peptide (SP), a key driver of post-mating changes (22–25). Ablation of main cells leads to failures in the induction of the main female post-mating responses: receptivity to remating remains high, and egg production unstimulated (26).

**Figure 1.**
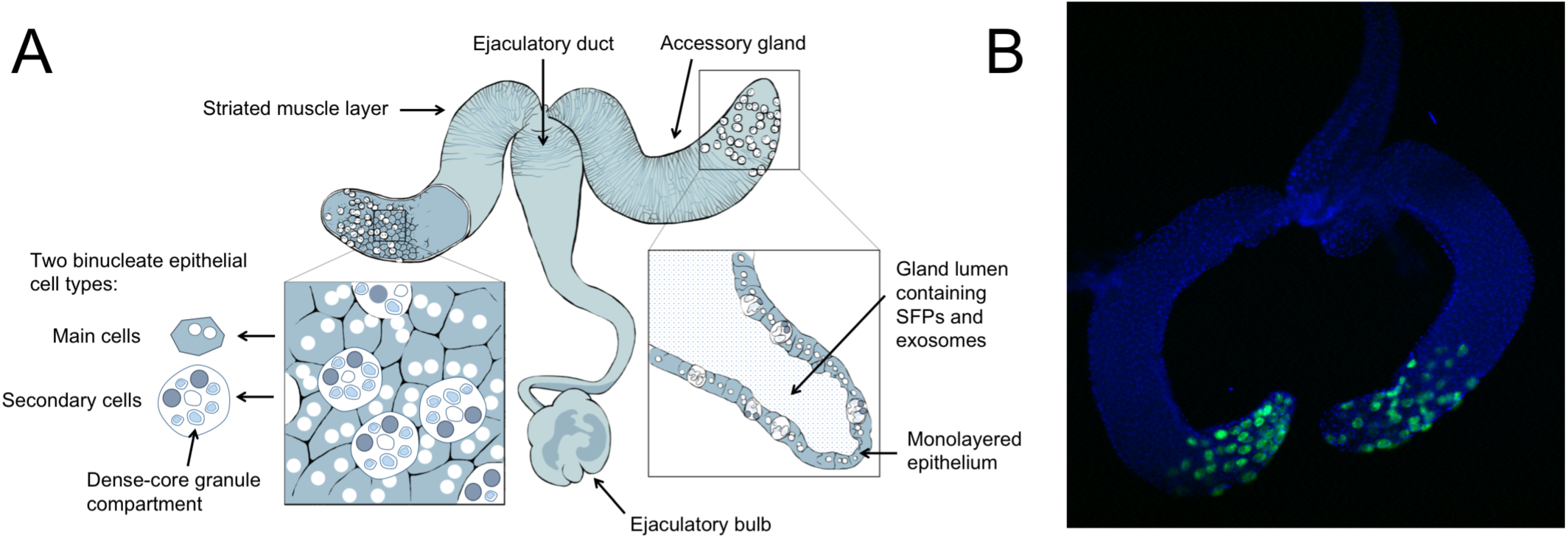
(A) The architecture of the *Drosophila melanogaster* male reproductive system. The testes, which branch off from where the two lobes of the accessory glands meet, are not shown. Figure adapted from (30). (B) Dissected glands from a control (*esg-GAL4* × *w^1118^*) male. Secondary cells fluorescence derives from *UAS-GFP_nls_*. Nuclei stained with DAPI. Image courtesy of Aashika Sekar.

The distal tips of each gland also contain a further subpopulation of ∼40 unusually large ‘secondary cells’ (20, 27; Fig. 1B). As with main cells, failures in secondary cell development are associated with defective post-mating responses: high receptivity, low fecundity (28, 29). This is partly attributable to glycosylation defects in ‘SP network’ proteins, which are required for the storage and gradual release of SP from the female sperm storage organs – the process through which SP’s effects are extended over several weeks (28). However, targeted suppression of BMP-signalling in adult secondary cells has more specific effects. While it suppresses the secretion of nanovesicles (‘exosomes’) and dense core granules – packages of secretory material that contain high concentrations of signalling molecules – it decouples female post-mating responses: fecundity is normally-stimulated, but sexual receptivity remains high (27, 30, 31). This raises the prospect that BMP-signalling in adult secondary cells acts as a highly-targeted mediator of reproductive processes. However, we do not know whether the phenotypic effects are restricted to those already identified, or whether secondary cell BMP-signalling is a potentially more global regulator of reproduction. This uncertainty also extends to the effects on the seminal proteome: does suppression of secretion by BMP-signalling inhibition in secondary cells cause highly specific changes to the seminal proteome or does it generate more extensive remodelling? In the present study, we use targeted suppression of BMP-signalling in adult secondary cells to test between these models at both the functional and proteomic level.

## RESULTS & DISCUSSION

### Sperm Storage is Compromised in Dad-Mated Females

We began by mating virgin females to males who possessed GFP-tagged sperm (32), and who overexpressed the transcriptional repressor of BMP-signalling Dad, which suppresses secondary cell secretion (31) (hereafter ‘Dad’ males), to test whether these secretions are required for normal sperm entry into storage. We found no significant difference in the number of sperm transferred (*F_1,53_*= 1.700, *p*=0.198; Fig. 2A), but the proportion that initially enter into the storage organs (seminal receptacle and paired spermathecae), and that are ultimately stored (5 hours post-mating; 32) was significantly lower in Dad-mated females (initial entry at 25 mins, *F_1,53_*= 5.340, *p*=0.024; Fig. 2B; 5hrs storage, *F_1,53_*= 5.043, *p*=0.029; Fig. 2C). This demonstrates a role for secondary cell activity in promoting normal sperm storage, which is surprising given that the number of offspring produced by Dad males has previously been shown to be normal (31). A potential mechanism for reduced storage in Dad-mated females is premature ejection of received sperm (33). However, we found no significant difference in the timing of ejection (*LRT*=0.892, *p*=0.345; SI Appendix, Fig. S1). Reduced sperm storage in Dad-mated females may instead be a consequence of loss of secondary-cell-derived exosomes, the prostate-derived equivalent of which in mammals are known to fuse with sperm and stimulate motility (34). Reduced storage could also arise if secondary cell BMP-signalling inhibition affected SFPs, such as the main cell-produced Acp36DE and/or its associated co-factors, which are known to collectively promote sperm storage (35–39).

**Figure 2.**
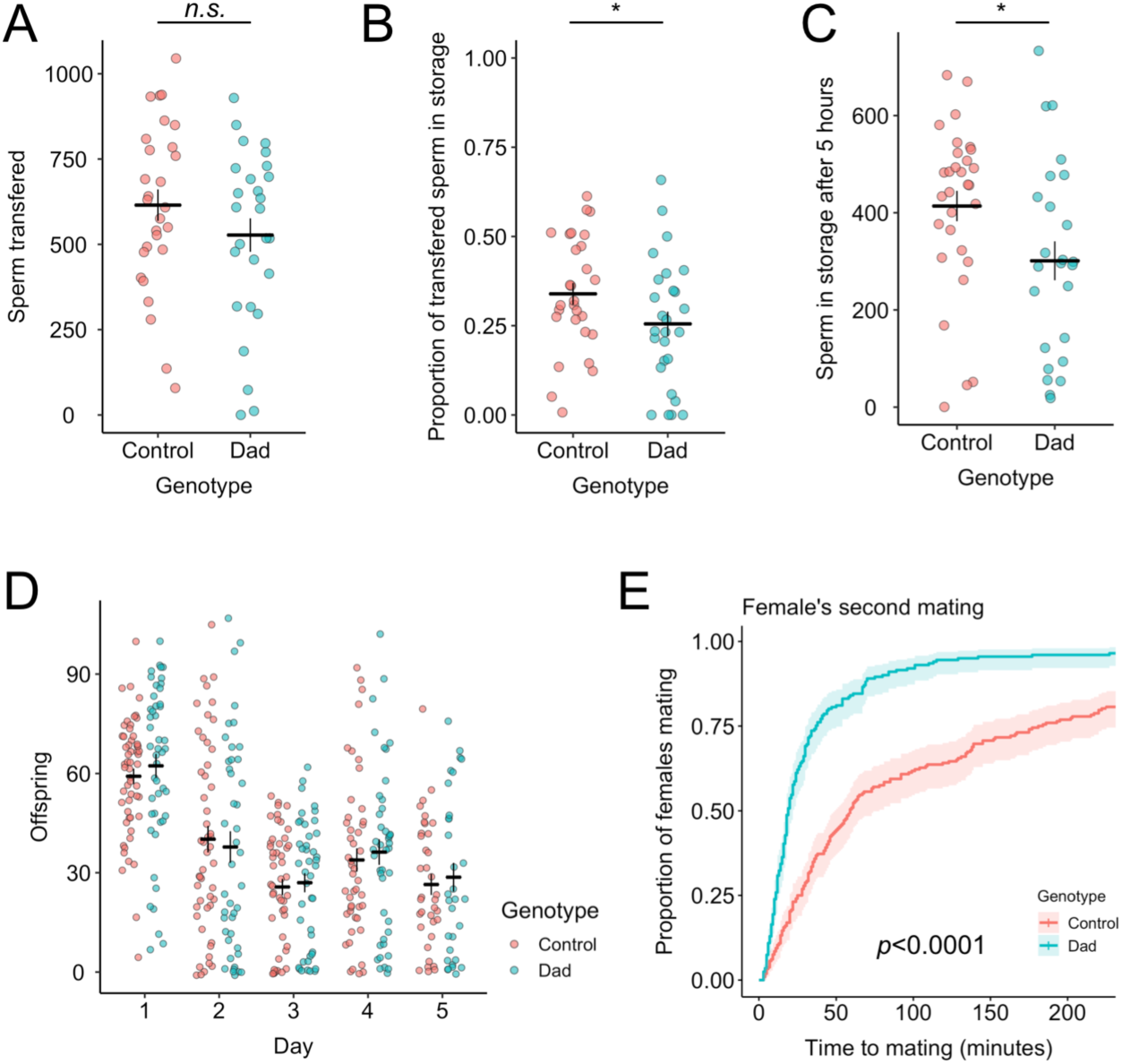
Defective sperm storage and decoupled post-mating responses in Dad-mated females. (*A*) The number of sperm present across all regions of the female reproductive tract 25 mins after the start of mating, *i.e.* the number transferred. n_Dad_=27, n_control_=28. (*B*) The proportion of transferred sperm that has entered into the storage organs (seminal receptacle and spermathecae) at 25 mins after the start of mating, n_Dad_=27, n_control_=28. (*C*) The number of sperm in storage at 5h after mating, n_Dad_=25, n_control_=30. (*D*) Daily offspring production, n_Dad_=47, n_control_=56. (*E*) The latency to remating by Dad- and control-mated females when presented with a second male 24h later, n_Dad_=276, n_control_=275. In *A-D*, horizontal bars represent the mean, with vertical bars representing ± 1 SE. Data are plotted with horizontal ‘jitter’. In *E*, confidence intervals are at 95%. * = *p*<0.05.

### Dad-Mated Females Show Decoupled Post-Mating Responses

Despite initially storing fewer sperm, we confirm previous work in finding that Dad-mated females show normal offspring production (31), additionally finding that this holds when females are far more fecund than in previous studies (likely due to the addition of live yeast to the fly food in our experiments, 40) and in both the short- and long-term (Genotype × Day, *F_4,346_*= 0.305, *p*=0.875; Genotype, *F_1,98_*= 0.007, *p*=0.932; Day, *F_4,346_*= 49.340, *p*<0.0001; Fig. 2D). We also confirm that Dad-mated females show abnormally high receptivity to remating (*LRT*=75.158, *p*<0.0001; Fig. 2E), an effect which is absent when flies are kept at low temperatures where Dad overexpression remains inactivated (see *Materials and Methods; LRT*=0.001, *p*=0.981; SI Appendix, Fig. S2), again supporting the finding that inhibition of BMP-signalling in secondary cells reduces male ability to induce refractoriness in their partners. This decoupling in the post-mating response is surprising given that both effects are driven by the binding of sex peptide (SP) to a specific receptor expressed in female reproductive tract neurons (41, 42). How these are mechanistically uncoupled remains unclear, but it may be that secondary cell secretions differentially affect interactions between SP and subpopulations of female reproductive tract neurons controlling receptivity (43, 44).

### Females Mated to Dad Males Over-Retain Sperm in the Seminal Receptacle Despite Normal Offspring Production

Because Dad-mated females store fewer sperm, but produce normal numbers of offspring, we predicted they would become sperm-depleted more rapidly. In contrast, we found significantly more sperm in the primary female sperm storage organ, the seminal receptacle, of Dad-mated females 7 days after copulation (*F_1,34_*= 12.568, *p* = 0.001; Fig. 3A). This effect was independent of the number of offspring produced (Genotype × Offspring, *F_1,33_*= 2.169, *p* = 0.150; Offspring, *F_1,34_*= 0.429, *p* = 0.517) and did not extend to the spermathecae, where we found no difference in sperm retention (*F_1,35_*=0.005, *p*=0.947; Fig. 3B). This result is only partially consistent with defective activity of SP: females that fail to receive SP are known to show defective release of stored sperm, as are females that receive a form of SP that cannot be cleaved from the sperm surface (45). However, defective SP activity causes a dramatic reduction in the rate of offspring production (28, 46), which is not exhibited by Dad-mated females. Moreover, defects in SP transfer and processing cannot explain the reduction in initial sperm storage in Dad-mated females as this process is known to be independent of SP (45). Thus, our data suggest both that (a) Dad-mated females show broad decoupling of post-mating responses (normal offspring production, but abnormal sperm release and receptivity) and, (b) the compromised ejaculate performance of Dad males is wide-ranging, affecting both SP-dependent (sperm release, receptivity) and SP-independent (sperm storage) reproductive processes.

**Figure 3.**
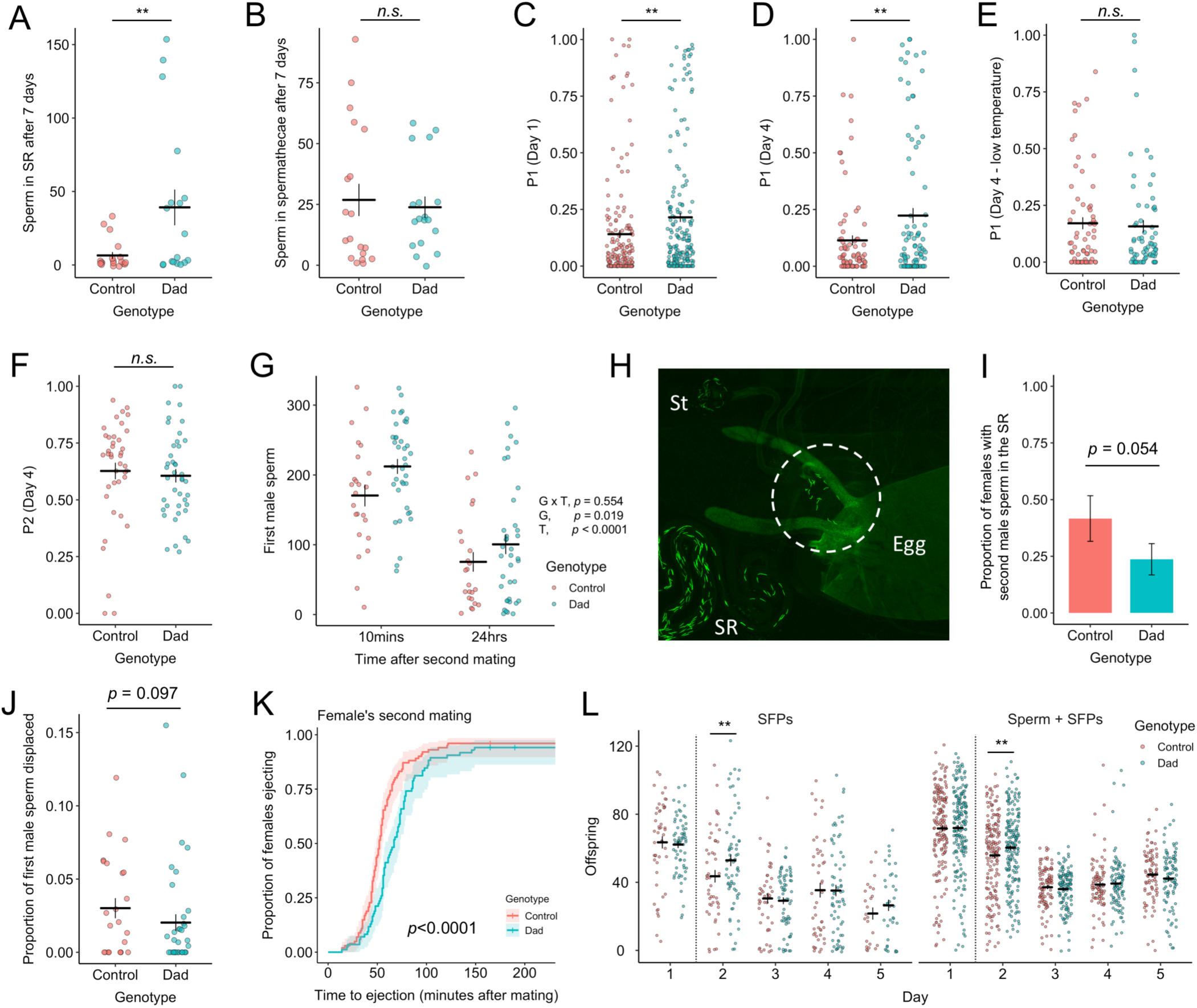
Dad-mated females over-retain sperm, provide higher first male paternity, and handle a second ejaculate differently. (*A*) The number of sperm in the seminal receptacle 7 days after singly mating to a Dad or control male. n_Dad_=18, n_control_=19. (*B*) As (*A*) but the total across both spermathecae. n_Dad_=18, n_control_=19. (*C*) First male paternity share when a female first mates to a Dad or control male and then a standardised competitor 24 hours later. Offspring collected over the 24h following remating. n_Dad_=190, n_control_=173. (*D*) As (*C*), but offspring collected in a 24-hour period 4 days after the female remated. n_Dad_=92, n_control_=81. (*E*) As (*D*), but conducted at 20°C to block Dad expression. n_Dad_=69, n_control_=67. (F) First male paternity share when a female first mated to a standardised competitor male and then a Dad or control male 24h later. Offspring collected over the 24h 4 days after remating. n_Dad_=43, n_control_=41. (*G*) Dad or control sperm across all regions of the female reproductive tract 10 minutes or 24 hours after remating to a standardised competitor. 10 mins: n_Dad_=38, n_control_=24; 24h: n_Dad_=38, n_control_=24. The *p*-values associated with Genotype (G), Timepoint (T), and their interaction in predicting sperm numbers are provided. (H) A female dissected at 5 hours after singly mating to a control male. Released sperm in the uterus are circled. SR, seminal receptacle; Sp, spermathecae. (I) Proportion of females where second male sperm has entered into the storage organs 10 mins after the start of mating. Females mated to a Dad or control male 24h previously. n_Dad_=38, n_control_=24. (J) As (I) but the proportion of the total first male sperm within the female reproductive tract that is found outside of the storage organs. n_Dad_=38, n_control_=24. (K) Latency to ejaculate ejection after previously Dad- or control-mated females remate with a standardised competitor. n_Dad_=85, n_control_=101. Confidence interval is 95%. (L) Daily offspring production by Dad- and control-mated females that secondarily mate to either a male transferring seminal fluid but no sperm or a normal second ejaculate. The dashed line gives the point at which the female remates. SFPs: n_Dad_=66, n_control_=48; SFPs + sperm: n_Dad_=193, n_control_=179. In panels A-G, I, J, and L, horizontal bars represent the mean, with vertical bars representing ± 1 SE of the mean or proportion. Data are plotted with horizontal ‘jitter’. * = *p*<0.05, **=*p*<0.01. Non-significant *p*-values between 0.05 and 0.1 are provided.

### Dad Males Acquire Higher Paternity Shares in Competitive Matings

*D. melanogaster* females can hold sperm from as many as 6 different males simultaneously (47). However, total female storage capacity is <1000 sperm, leading to sperm competition between rival males (32). Consequently, males are presumed to be under selection to both displace resident sperm from storage when mating with non-virgin females (‘offensive sperm competition’) and in turn, to produce sperm that resist displacement by incoming ejaculates (‘defensive sperm competition’) (48). To test whether these abilities are mediated by secondary cells, we first mated a Dad or control male to a virgin female, who then remated 24hrs later with a standard male competitor. Both the females and competitor males carried a recessive *sparkling* (*spa*) eye marker, which allowed us to assign paternity of the resulting offspring (49–52). We found that Dad males gained significantly higher first-male paternity shares (‘P1’) in offspring produced over the first day after female remating (*F_1,360_*= 9.445, *p*=0.002; Fig. 3C). This effect was still present in offspring produced in 24-hour periods at day 4 (*F_1,171_*= 11.525, *p*=0.009; Fig. 3D) and day 6 (*F_1,105_*= 7.424, *p*=0.008) after the female remated. It was also independent of remating latency either overall (*F_1,359_*= 0.264, *p* = 0.608; SI Appendix, Fig. S3) or as an interaction with male genotype (*F_1,357_*= 0.329, *p* = 0.567), which suggests that the elevated P1 of Dad males is not an artefact arising through a lack of remating by control-mated females. No P1 differences were detected when flies are kept at low temperatures where Dad overexpression remains inactivated (day 1, *F_1,134_*= 1.717, *p*=0.192; day 4, *F_1,131_*= 1.027, *p*=0.313; Fig. 3E), confirming that the effect is caused by inhibition of BMP-signalling in secondary cells. Next, we reversed the mating order, such that Dad or control males mated to a female previously mated to a *spa* male, and found no effect on paternity share (P2; 24 hours, *F_1,81_*= 0.246, *p*=0.621; 4 days, *F_1,80_*= 1.814, *p*=0.182, Fig. 3F). Thus, the effect of secondary cell secretions on sperm competition performance are mating-order specific.

### Over-Retention of Dad Sperm Provides a Mechanism for Enhanced Paternity Share

Under single-mating conditions, Dad-mated females retain more sperm 7-days after mating (Fig. 3A). Under double-mating conditions, Dad males achieve higher paternity shares (Fig. 3C,D). Thus, a possible mechanism for the increased paternity share is Dad-mated females having greater numbers of first male sperm in storage at the time of second mating compared to control-mated females. This mechanism would explain why we detect no differences in P2 and would be partially consistent with previous work on failure in secondary cell development, which showed over-retention of sperm and improved paternity share, but crucially alongside dramatically-reduced offspring production (28). However, given that Dad-mated females initially store fewer sperm (Fig. 2C) and display normal productivity (Fig. 2D) we predicted a different mechanism: that the elevated paternity share achieved by Dad males acts through enhanced resistance to displacement by a second male ejaculate. To test this, we counted sperm across all regions of the female reproductive tract at two time-points after the start of a female’s second mating: 10 minutes (∼halfway through mating) and 24 hours. By selecting these time-points, we were able to ask whether the P1 advantage in Dad-mated females is present from the outset of a female’s second mating (*i.e.* Dad-mated females have retained more sperm) or whether it develops over the course of second male sperm entering into storage.

Overall, we found significantly higher quantities of first male sperm throughout the female reproductive tract (in storage or displaced into the uterus; *F_1,120_*=5.616, *p*=0.019; Fig. 3G) in Dad-mated females. This effect was independent of the time-point after mating (Genotype × Timepoint, *F_1,119_*=0.351, *p*=0.554; Fig. 3G), but contrary to our prediction, there was a trend for the degree of difference between Dad and control sperm number to be diminished 24 hours after re-mating. Thus, the P1 sperm advantage in Dad-mated females appears to be present at the start of a female’s second mating and, if anything, remating appears to weaken, not reinforce the sperm advantage of the Dad male. This also means that despite Dad-mated females initially storing reduced quantities of sperm (Fig. 2C), they hold more in storage relative to control-mated females by the time of their second mating (Fig. 3G). Greater retention of sperm is a known consequence of SP dysregulation, but in these cases it is partly explained by females using fewer sperm because they produce fewer offspring (28, 45). Why, then, does reduced sperm release in Dad-mated females not translate into reduced offspring output (Fig. 2D)? The most parsimonious explanation is that Dad-mated females achieve the same number of fertilisations as control-mated females, but release fewer sperm per fertilisation. Previous estimates suggest that females release 1-5 sperm per fertilisation, but are able to modulate the efficiency of sperm use in response to variation in environmental quality (reviewed in 53). While sperm use is challenging to measure directly, on the rare occasions where we found eggs in the uterus of dissected females we did find instances where large number of sperm (up to 17) were associated with an egg (Fig. 3H), suggesting that sperm use may be more inefficient than previously suggested. This inefficiency may be particularly pronounced when the storage organs are largely full, as would be the case so soon after mating (5 hours). Despite appearing wasteful, profligacy in sperm release may be adaptive if it encourages further competition between sperm of varying quality, with consequences for offspring fitness (54–56).

### Altered Dynamics of Second Male Ejaculates in Dad-Mated Females

Dad-mated females treat potential sexual partners differently by showing higher receptivity to remating. We therefore sought to test whether they treat second male sperm differently. We first looked at the rate at which second male sperm are stored. It is already known that if a male fails to transfer Acp36DE both his sperm and those transferred by the next male show compromised storage, despite the second male presumably transferring Acp36DE himself (10). Dissecting females 10 minutes after starting a second mating, we found a non-significant trend for slowed entry of second male sperm in previously Dad-mated females (*F_1,59_*= 3.718, *p* = 0.054; Fig. 3I) and reduced displacement of first male sperm at this time point (first male sperm in the uterus/total first male sperm across all regions of the reproductive tract; *F_1,61_*= 2.836, *p* = 0.097; Fig. 3J).

We next tested for differences in the timing of female ejection. The length of time a female retains a second male ejaculate after remating influences the outcome of sperm competition: the longer it takes a female to eject, the greater the opportunity for second male sperm to enter into storage and displace resident sperm (57). We therefore predicted that Dad-mated females would eject sperm earlier, thereby terminating the displacement of first male sperm, and promoting the paternity share advantage experienced by Dad males (Fig. 3C). Contrary to expectation, Dad-mated females were significantly slower to eject after their second mating (*LRT*=17.981, *p*<0.0001; Fig. 3K). This should weaken the advantage experienced by Dad males that arises through over-retention of sperm by their female partners. Indeed, this weakening could explain the slight decrease in the degree of difference between Dad and control sperm number in the 24 hours after re-mating relative to 10 minutes after re-mating (Fig. 3G). Ultimately, this result suggests that female treatment of a second ejaculate is influenced by features of the first male’s ejaculate.

Finally, we tested whether offspring production after a second mating differs depending on whether a female first mated with a Dad male or a control. As second males we used either males transferring both sperm and seminal fluid or spermless *son-of-Tudor* males that transfer seminal fluid but no sperm. This allowed us to identify the relative importance of second male sperm and seminal fluid in driving any detected effects. We found a significant interaction between day since mating and first male genotype on daily offspring production (*F_4,1432_*=2.740, *p*=0.027; Fig. 3L). This appears to be driven by a short-term increase in offspring production by Dad-mated females exclusively in the 24 hours following remating (*t ratio=*2.663, *p*=0.008). This effect was independent of whether the female received second male sperm (First male × Second male × Day, *F_4,1398_*=0.577, *p*=0.679; First male × Second male, *F_1,400_*=0.096, *p*=0.757), demonstrating that it is specifically attributable to the second male’s seminal fluid. A potential mechanism for this short-term boost in offspring production in Dad-mated females is second males transferring larger quantities of fecundity-stimulating SFPs when mating with Dad-mated females compared to those females previously mated to controls. There is good precedent for this: males strategically decrease their transfer of the short-term acting, fecundity-stimulating SFP ovulin when they detect that they are mating with a mated female (58). Given the high receptivity of Dad-mated females, second males may perceive them as virgins and transfer higher quantities of SFPs such as ovulin, though this remains to be tested.

### The SFP Proteome is Remodelled in Dad Males

The phenotypic effects we find in Dad-mated females are likely to arise through changes to the production, transfer, and protein composition of seminal fluid, particularly given that BMP-signalling promotes secondary cell secretion (27, 30). This change may operate exclusively through secondary cells or, if there is cross-talk between cell-types, also via their influence on main cells. To this end, we performed label-free quantitative proteomics on the accessory glands of Dad and control males dissected either before or immediately after mating. This pre- and post-mating approach has previously been shown to provide a deep analysis of the seminal proteome, sensitive to low abundance proteins, while exposing patterns of differential SFP production, depletion, and transfer (19, 51). We detected 1194 proteins on the basis of at least 2 unique peptides (as in 19, 59), of which 88 are SFPs known to be transferred to females (see *Materials and Methods*). A principal component analysis (PCA) conducted on these 88 SFPs showed full separation of samples in relation to both genotype and mating status (Fig. 4B). Analysis of the extracted scores showed that PC1, which described the majority of variance (60.8%), was associated with the interaction between mating and genotype (Table S1). PC2 was significantly described by male genotype and captures an axis of variation (7.8%) associated with divergent responses among SFPs in the extent to which their abundance was affected by secondary cell disruption. Thus, as expected, inhibition of BMP-signalling in secondary cells changes the SFP composition of the accessory glands.

**Figure 4.**
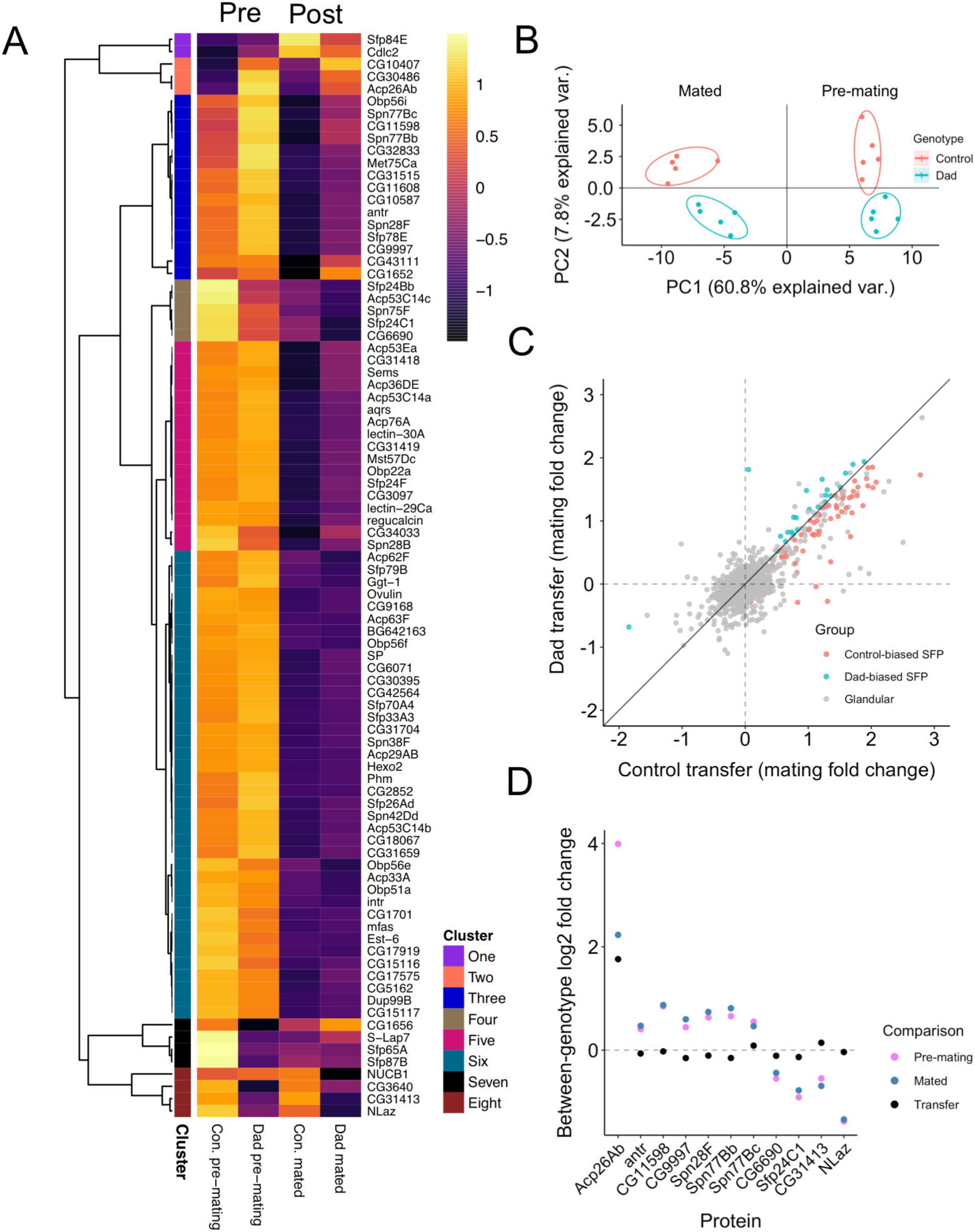
Quantitative proteomics reveals defects in SFP transfer in Dad males. (*A*) A heatmap showing the abundance patterns of SFPs. Columns 1 and 2: males dissected prior to mating; columns 3 and 4: males dissected 25 minutes after mating. Columns 1 and 3: Dad males; Columns 2 and 4: control males. Row annotations highlight membership of higher-order clusters based on a Pearson correlation distance metric. (*B*) Output of a PCA conducted on abundances of the 88 detected SFPs. Points coloured according to male genotype. Mated glands on the left, pre-mating glands on the right of *x*=0 line. Ellipses denote 80% normal probability. (*C*) Correlation between Dad and control pre- vs post-mating fold changes (degree of transfer) for each SFP. Red gives SFPs transferred in greater quantities by control males, blue gives SFPs transferred in greater quantities by Dad males. Grey denotes non-SFPs. (*D*) Log_2_ fold changes for three different between-genotype comparisons for each of 11 SFPs identified as showing a significant abundance change in response to BMP-signalling suppression. Comparisons: pre-mating (pink), post-mating (blue), and transfer to females (black). Positive values indicate greater abundance in Dads.

### Split Responses of the Seminal Proteome to Suppression of Secondary Cell BMP-Signalling

To test for patterns among SFPs in their response to BMP-signalling suppression in secondary cells, we undertook a hierarchical clustering analysis across genotypes and mating treatments (Fig. 4A). Responses of SFPs to genotype appear variable with multiple higher-order clusters identified. The changes did not suggest a complete loss of any SFPs in Dad males. Instead, we find evidence of quantitative changes in the abundance of some SFPs. Indeed, we find that a majority of SFPs are transferred in smaller quantities in Dad males compared to controls (67% of SFPs show smaller change in Dad; 2-tailed binomial test, *p*=0.002; Fig. 4C). Following false detection rate (FDR) correction, we failed to identify any SFPs showing the significant mating × genotype interaction that would indicate high-confidence differences in transfer. This may in part due to low power (5 samples per treatment combination), but it could also be due to any differences in transfer being relatively small, which seems to be the case for most SFPs (Fig. 4C). However, we found that 11 of the 88 SFPs show a significant response to genotype (Fig. 4D; Table S2; Fig. S4). This list did not include SP or Acp36DE, two candidate proteins that could be influencing the receptivity (Fig. 2E) and sperm storage (Fig. 2C) phenotypes, respectively, that we detect in Dad-mated females. A further 26 differentially abundant glandular proteins (*i.e.* non-SFPs) are given in Table S3. Thus, while SFPs make up just 7.4% of the proteins we detect (88/1194), they make up 29.7% (11/37) of the proteins showing a significant difference in abundance in Dad males, suggesting a disproportionate effect of BMP-signalling suppression on the seminal fluid proteome.

7 of the 11 differentially abundant SFPs showed higher abundance in Dad glands (Acp26Ab, antr, CG11598, CG9997, Spn28F, Spn77Bb, Spn77Bc), 4 showed higher abundance in control glands (CG6690, Sfp24C1, CG31413, NLaz). CG9997 is thought to be specifically expressed in secondary cells, but we did not find significant differences in abundance in other SFPs thought to be exclusively produced in the secondary cells, such as CG1652, CG1656, and CG17575 (28). Therefore, suppression of BMP-signalling does not appear to block production of these secondary cell proteins, and its effects on their abundance seem to be selective.

Acp26Ab stands out from the other differentially abundant SFPs in the scale of its expression differences: 16x more abundant in Dad pre-mating glands and 8x more abundant in Dad mated glands. This suggests, counterintuitively, that Dad males increase the transfer of this SFP. Consistent with this, Acp26Ab had the lowest FDR-corrected genotype × mating *p*-value of the 1194 proteins we tested (*p*=0.059). Interestingly, previous work has shown that Acp26Ab is present in both main and secondary cells within the first day of eclosion, but after 5 days is only present within the dense core granules of secondary cells (60), a pattern that suggests Acp26Ab is produced by main cells and trafficked to secondary cells. Suppression of BMP-signalling in secondary cells may disrupt this process of inter-cellular transport and lead to over-production of Acp26Ab by main cells. Similarly, CG11598 has previously been shown to be present in both main and secondary cells. In a previous transcriptomic study, manipulation of secondary cell development led to a large downregulation of *CG11598* expression, the magnitude of which was suggested to only be accountable for by changes in main cell activity (21). Surprisingly, we find that the abundance of CG11598 changed in the opposite direction, being more abundant following suppression of secondary cell BMP-signalling. Collectively, the changes we detect in Acp26Ab and CG11598 suggest a role for the secondary cells in mediating the activity of main cells, perhaps via cell-cell signalling.

In 9 of 11 of these proteins, the between-genotype fold change became more Dad-biased after mating (blue dot above pink dot, Fig. 4D). Indeed, looking across all 88 SFPs we find that the majority of SFPs are at higher abundance in Dad glands prior to mating (65%, 57/88; 2-tailed binomial test, *p*=0.007) with the number increasing after mating (73%, 64/88; 2-tailed binomial test, *p*<0.0001). We offer two explanations for why the majority of SFPs are initially at higher abundance in Dad males. Firstly, Dad males may overproduce SFPs, perhaps due to disruption to main cell/secondary cell signalling. Secondly, if males suffer even slightly reduced SFP transfer in each mating then they may accumulate over-retained SFPs following the previous day’s triple-matings, which we provided to clear the glands of products produced prior to expressing Dad (*Materials and Methods*; as in (27, 31)). In either case, the differences in transfer for the significantly differentially abundant SFPs are surprisingly small given the clear between-genotype differences in their abundance within the gland (Fig. 4D). This suggests that there may be mechanisms that regulate the quantity of accessory gland secretion that is transferred to females independently of both the quantity within the gland and secondary cell activity.

## CONCLUSIONS

We conclude that BMP-signalling in adult secondary cells is a major mediator of manifold reproductive processes. These findings have broad implications for our understanding of how ejaculates evolve. Firstly, ejaculate evolution may be constrained. Although normal secondary cell activity inhibits male defensive sperm competition performance, it is required to reduce female receptivity to remating. Given that the latter ability is the wild-type condition, it seems likely that the benefits loss of secondary cell secretion brings to paternity share are outweighed by the benefits of suppressing female receptivity to remating. However, the question remains why males apparently aren’t able to simultaneously maximise performance in both. Such intra-ejaculate trade-offs in function may represent an under-appreciated constraining force on ejaculate evolution. Secondly, our data demonstrate that the composition and function of the ejaculate depends on the integrated activity of the two constituent cell-types of the accessory glands. Thus, evolutionary changes to the architecture of seminal fluid-producing tissues would have knock-on consequences for ejaculate composition and function. Interestingly, secondary cell number is variable between *Drosophila* species – they have even been lost entirely in *Drosophila grimshawi* (15). In light of our results, we would predict covariance between accessory gland cellular architecture and variable aspects of mating biology, such as mating rate and sperm competition intensity, across the Drosophila phylogeny. Given that we find an element of modularity in ejaculate design, with normal offspring production being exclusively driven by main cell activity in adults, it may be that some reproductive functions are insulated from changes in a given part of the male reproductive system. Ultimately, by taking an evo-devo approach to male reproductive tissues we may begin to understand how ejaculate function and composition evolve.

## MATERIALS AND METHODS

### Fly Stocks and Husbandry

Males with disrupted secondary cell secretion were generated by crossing esg^ts^ F/O flies (genotype: *w; esg-GAL4 tub-GAL80^ts^ UAS-FLP/CyO; UAS-GFP_nls_ actin>FRT>CD2>FRT>GAL4/TM6)* to *w^1118^* flies into which a *UAS-Dad* transgene had been backcrossed (‘Dad’ males)(27, 31). For controls, we crossed esg^ts^ F/O to flies from a *w^1118^* background (‘control’ males). The *esg-GAL4* system incorporates a temperature-sensitive GAL80, which inhibits GAL4 and suppresses the activation of *Dad* expression below 28.5°C (see 31). Where sperm counts were undertaken, we backcrossed the *GFP-ProtB* construct, which labels the heads of sperm (32), into our Dad and *w^1118^* lines for 6 generations. All females were from a Dahomey wild-type background into which the *spa^pol^* recessive eye-marker had previously been backcrossed for 4 generations. All competitor males were of this same genotype or, where sperm counts were conducted, this genotype carrying a *RFP-ProtB* construct (32).

All flies were reared at standardised larval densities of ∼200 in 250mL bottles containing 50mL of Lewis medium (as in 61). Larvae were left to develop at a non-permissive temperature of 20°C on a 12:12 L:D cycle. Upon eclosion, we collected males under ice anaesthesia and separated them into groups of 8 to 12 in 36mL Lewis medium-containing plastic vials, supplemented with *ad libitum* yeast granules. To activate the expression of *Dad* (where present), we immediately moved these vials to 30°C where they remained for the full duration of experiments. To verify that phenotypes were specifically attributable to *Dad* expression, we repeated some experiments at a non-permissive temperature of 20°C. In these experiments, flies were moved to 20°C after eclosion where they remained for the full duration of experiments. The day before using Dad or control males, each was mated to three virgin females to deplete, as much as possible, the accessory gland lumen of any secondary cell products produced before activation of the *Dad* transgene. We delivered a single female at a time, removing the female after mating. Following the end of the third mating, we moved the male to a fresh, yeast-supplemented vial.

The rearing, collection, and grouping of flies from all other lines was performed following the methods outlined above. However, in these cases rearing was conducted at 25°C with us moving flies to 30°C the evening before use in experiments. We reared all flies and performed all experiments in controlled-temperature rooms on 12:12 light:dark cycles. All flies were between 3 and 5 days old at the time of first experimental mating.

### Sperm Count Experiments

We conducted the initial sperm transfer experiment in two blocks. Females were frozen at 25 minutes or 5 hours after the start of mating (ASM). We conducted the post-first-mating retention experiment in one block. Here, females were frozen 7 days after mating. We conducted the post-second-mating sperm dynamics experiment in two blocks. Here, females were frozen at 10 minutes or 24 hours after second mating. Females in all experiments were randomly assigned a freezing time-point prior to mating. Offspring were collected and counted between mating and freezing where appropriate. Females were flash-frozen in liquid nitrogen and stored at −80°C until dissection, which we performed under light microscope in PBS. We retained the female reproductive tract from the vulva through to the common oviduct, sealed the slides using (Fixogum, Marabu), and stored slides at 5°C. We imaged the slides using a Zeiss 880 confocal microscope and processed the images by taking an average intensity Z-projection in the Fiji distribution of ImageJ (62) to condense Z-stacks into a single image for easier counting. We manually counted sperm using the multi-point tool in Fiji. We performed all dissections and sperm counts blind to treatment. We omitted any samples that showed no GFP sperm due to the possibility of heterozygosity for the *GFP-ProtB* chromosome in our stock populations.

### Sperm competition outcome and post-mating response assays

For P1 defensive sperm competition assays, we aspirated single Dad or control males into yeasted vials containing an individual virgin *spa^pol^* female. We monitored all matings, recording the time males were introduced, mating began, and when mating finished. From these data we calculated the duration of and latency to mating. After mating, we disposed of the males and left the females to oviposit. The following morning, we individually aspirated mated females into a yeasted vial containing a pair of *spa^pol^* males, grouped under ice anaesthesia the previous day. Again, we monitored all matings and recorded duration and latency. We introduced females in the order they had finished mating the previous day. Previous work has shown that Dad-mated females remain highly receptive to remating (31), so we staggered the introduction of Dad- mated females to minimise any systematic difference between treatments in inter-mating interval. Following the end of mating, we discarded the two males and moved the females to 25°C, transferring them into a fresh, yeasted vial every 24 hours. We allowed the resulting progeny to develop, freezing the vials after the adults eclosed. We then counted offspring and scored their eye phenotype in order to assign paternity. By adopting this same approach but reversing the mating order, we tested for an association with offensive sperm competition performance (P2). We performed three blocks of a repeat of the P1 experiment conducted entirely at a non-permissive temperature of 20°C. We obtained P1 data across 6 experimental blocks at 30°C. In each of these, we collected offspring for at least 24 hours after the female’s second mating. In one replicate, we collected offspring for 6 days to test for the persistence of any detected differences. Within four of these replicates, we varied the identity of the second mating male. Here, prior to first mating to a Dad or control male, females were randomly assigned (a) no second mating, (b) a *spa^pol^* second mating, or (c) a spermless, *son-of-Tudor* mating. In these variants, we collected offspring over four days after second mating to gain additional information relating to short- and longer-term patterns of offspring production.

### Female ejection assays

We followed the P1 experimental setup outlined in the preceding section, but moved females to 3D-printed, black plastic chambers immediately after a first or second mating. These chambers, of printing resolution 0.2mm, were cuboids of 34mm × 33mm × 9mm with a half-sphere concavity of dimensions 20mm × 20mm × 7mm. A .stl file of this design is included as a supplementary file for use by other researchers. We used a glass coverslip to cover the concavity once a female had been introduced. We checked each chamber for the presence of an ejected sperm mass every 10 minutes under a light microscope. We ran this experiment four times: twice for each of the females first (Dad or control) and second (*spa^pol^*) mating.

### Proteomics experiment

We randomly assigned males a mating treatment (‘pre-mating’ or ‘mated’) and paired within a genotype. We aspirated the ‘mated’ treatment male within each pair into a yeasted vial containing an individually isolated 4/5-day old virgin female. At this same point, the ‘pre-mating’ male from the pair was introduced to an empty, yeasted vial. We flash-froze ‘mated’ males in liquid nitrogen 25 minutes after the start of mating, freezing their ‘pre-mating’ partner at the same time. This paired freezing approach ensures that the distribution of freezing times is equivalent between mated and pre-mating males. Frozen males were stored at −80°C until dissection.

For each sample, we pooled 20 pairs of accessory glands, which we dissected under a light microscope on ice in a drop of ice-cold PBS. We took care to remove the seminal vesicles and testes, and severed the glands from the distal end of the ejaculatory duct. Dissected glands were then transferred to an Eppendorf tube containing 25µl of PBS, which we stored at −80°C. In total, we had 20 samples: five for each of the four treatment permutations (mated, Dad; pre-mating, Dad; mated, control; pre-mating, control). We ran this experiment five times in order to produce five independent biological replicates. Our quantitative proteomics analysis was conducted in accordance with the gel-aided sample preparation (GASP) protocol outlined in detail elsewhere (19, 63). Details of this method, the LC-MS/MS platform, and the data processing and normalization are given in *SI Materials and Methods*.

The mass spectrometry proteomics data will be deposited to the ProteomeXchange Consortium via the PRIDE (64) partner repository.

### Statistical analysis

We conducted all analyses with R statistical software (version 3.5.1)(65) in RStudio (version 1.1.456)(66). We assessed the significance of variables in linear and generalized linear models by dropping individual terms from the full model using the ‘drop1’ function. Where the interaction term was non-significant we refitted the model without it. We determined model fit by visual inspection of diagnostic plots (67). Where multiple measurements were taken from the same female, as in analyses of day-by-day female offspring production, we used linear mixed effects models that accounted for female identity as a random effect. In our day-by-day analysis of female offspring production, our starting model contained a three-way interaction (male 1 × male 2 × day) along with two random effects (block and female ID). We used a stepwise algorithm (‘step’ function) to identify the best model by AIC. Associated *p*-values were generated using Satterthwaite’s method (68). To analyse latency to mating and ejection, we ran Cox proportional hazard models using the *survival* package (69, 70) and graphed the results using ‘ggsurvplot’ in the *survminer* package (71). We analysed proportional data, relevant for paternity shares (P1 and P2) and some sperm count data, using generalised linear models. In all cases, we used a quasibinomial extension to account for the overdispersion we detected. When analysing the number of sperm retained in the seminal receptacle after 7 days, we used a quasipoisson distribution to correct for overdispersion. We limited all analyses to matings lasting longer than 7 minutes and which gave rise to fertile offspring to exclude disturbed or pseudo-matings (72). In our analysis of first male sperm retention after a second mating, we winsorized one extreme significant outlier (as determined by two-tailed Grubbs’ test) found to exert disproportionate leverage in our models (73).

Our assessment of whether a protein was a SFP was based on a reference list provided by Mariana Wolfner (Cornell University, NY) and Geoff Findlay (College of the Holy Cross, MA) and updated to include the high confidence SFPs from Sepil *et al.* (19). We also included Intrepid (intr), despite it not having been conclusively shown to be transferred to females, as we find it at significantly lower abundance in mated glands and because it is known to function in the sex peptide network (16). All analyses were performed on log_2_ transformed values to standardise the variance across the dynamic range of protein abundances. Fold changes were calculated using per-treatment means (taken across the five replicates). Our hierarchical clustering analysis was conducted on the mean per-SFP abundance taken across the five replicates for each treatment permutation and used a Pearson correlation distance metric. We plotted the results using the *pheatmap* package (74). We conducted a PCA on SFPs using the ‘prncomp’ function in *stats*. Variables were scaled to have unit variance and shifted to be zero-centred. We ran linear models on the PC scores to test for associations between PCs and our variables. For our differential abundance analysis, we iterated a linear model over all detected proteins across the 20 samples, including genotype, replicate, and mating status as factors. We used a tail-based false discovery rate correction from the *fdrtool* package (75).

## ACKNOWLEDGEMENTS

We thank Josephine Hellberg and Aashika Sekar for images, Mariana Wolfner and Geoff Findlay for sharing their list of SFPs, and Stefan Lüpold for providing the *GFP-ProtB* line. Thanks also to Natasha Gillies, Rebecca Dean, and Lynn Marie Johnson for advice on the statistical analysis and to Alex Majane, Artyom Kopp, and David Begun for drawing our attention to the absence of secondary cells in *D. grimshawi*. This work was funded by the EP Abraham Cephalosporin-Oxford Graduate Scholarship to B.R.H., with additional support from the BBSRC DTP. S.B., P.C., R.F., and B.M.K. were supported by the Wellcome Trust (097813/11/Z) and John Fell Fund (133/075). C.W. was supported by the BBSRC (BB/N016300/1, BB/R004862/1) and CRUK (C19591/A19076). I.S. and S.W. were supported by a BBSRC fellowship to S.W. (BB/K014544/1).

## Notes

The authors declare no conflicts of interest

