## Supplementary information appendix for "BMP-signalling inhibition in *Drosophila* secondary cells remodels the seminal proteome, and self and rival ejaculate functions"

**This PDF file includes:**

Supplementary information text
References for SI
Figs. S1 to S4
Table S1 to S3

### Supplementary Information Text

### Materials and Methods

#### Proteomics sample preparation

We prepared our samples for proteomic analysis in line with the previously published GASP protocol (1,2). The glandular tissue was first macerated using a clean pestle for a timed interval of one minute. To lyse the cells, we added 25µl of Pierce RIPA (Radioimmunoprecipitation Assay) Buffer, which we dripped over the pestle to flush residual tissue back into the sample. Next, we incubated the lysate with 50mM of the reducing agent DTT (Dithiothreitol) for approximately 10 to 20 minutes. To this, we added at room temperature an equal volume of 40% acrylamide/Bis solution (37.5:1 National Diagnostics) to facilitate cysteine alkylation to propionamide. Next, we added 5µl of 10% APS (ammonium persulphate) and an equivalent quantity of TEMED (tetramethylethylenediamine) to trigger the polymerization of acrylamide and form a gel plug. This plug was subsequently shredded via centrifugation through a membrane-less Spin-X filter insert (CLS9301, Sigma/Corning). Gel-fragments from this process were fixed in 40% ethanol/5% acetic acid before washing with a solution of 50mM ammonium bicarbonate, 1.5M Urea, and 0.5M Thiourea, which was then removed with acetonitrile. 250µl of dilute trypsin (Promega) was next added and the solution left at 37°C overnight to promote digestion of the immobilised peptides. The resulting peptides were extracted via two repeated ACN (acetonitrile) replacements, dried, desalted in Sola SPE columns (Thermo), and then suspended in 0.1% FA (formic acid), 2% ACN before LC-MS/MS (liquid chromatography-mass spectrometry/mass spectrometry) analysis.

**LC-MS/MS.** For peptide analysis, we used a LC-MS/MS platform composed of a Dionex Ultimate 3000 and a Q-Exactive mass spectrometer (Thermo). Peptide loading took place in a solution of 0.1% TFA (trifluoroacetic acid) in 2% ACN on a trap column (PepMAP C18, 300µm x 5m, 5µm particle, Thermo). For separation, we used an easy spray column (PepMAP C18, 75µm x 500m, 2µm particle, Thermo) with a gradient 2% ACN to 35% ACN in 0.1% FA in 5% DMSO (dimethyl sulphoxide). For MS spectra collection, we used a resolution of 70,000 in profile mode on the Q-Exactive (ion target = $3x10^6$). We selected the top 15 most intense features selected for subsequent MS/MS analysis (resolution of

17,500). The following parameters were set: dynamic exclusion = 27 seconds; AGC target = $1x10^5$; isolation width = 1.6m/z; and maximum acquisition time = 100ms.

**MS data processing.** The following MS data processing pipeline has previously been outlined in Sepil et al. (2). We imported the RAW data into Progenesis QIP (version 4.1.6675.48614), exporting spectra as MGF files using the 200 most intense peaks without deconvolution for searching. Peptide identification used the D. melanogaster UniProt reference proteome as a search target, with database retrieval conducted on 27/09/2017 (23306 sequences) in Mascot 2.5.1. Search parameters were set as follows: Oxidation (M), Propionamide (K), and Deamidation (N,Q) as variable modifications; Propionamide (C) as a fixed modification; one missed cleavage site; 0.05 Da fragment mass accuracy; 10ppm precursor accuracy. Prior to importing the search results into Progenesis for quantification via the Top3 method, we applied a peptide-level 1% FDR alongside a further Mascot ion cut-off of 20. The resulting protein abundance data was subsequently normalised using the internal Progenesis algorithm to a set of housekeeping proteins.

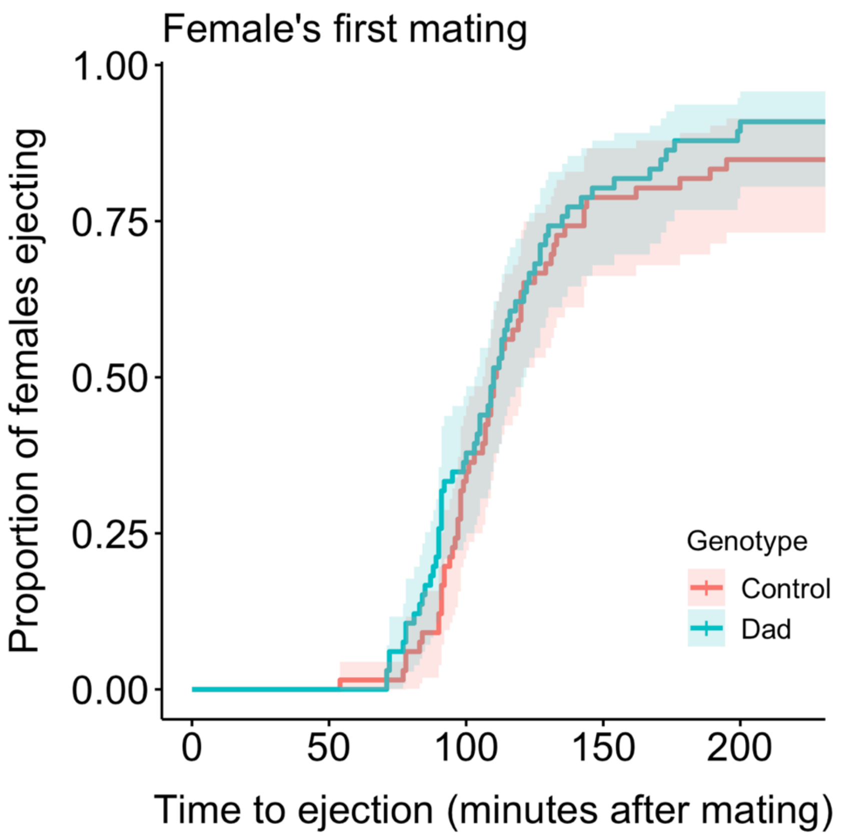


**Figure S1.** No significant difference in the timing to ejaculate ejection after a female singly mates with either a Dad or control male. $n_{Dad}$=66, $n_{Control}$=66. Confidence intervals are at 95%. Data pooled from two experimental blocks. $p$=0.345.

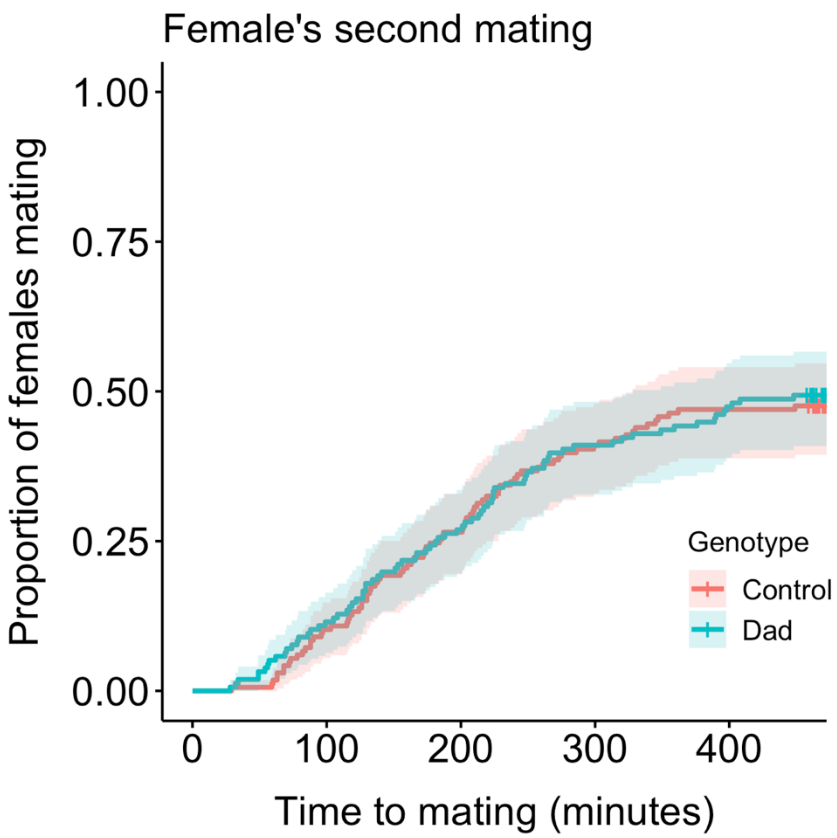


**Figure S2.** No difference in the latency to remating after previously Dad- or control-mated females are provided with a second mating opportunity 24 hours later. Here, both Dad and control males were reared at temperature of 20°C where Dad is not expressed. $n_{Dad}$=156, $n_{Control}$=166. Confidence intervals drawn at 95%. Data pooled from three experimental blocks. $p$=0.981.

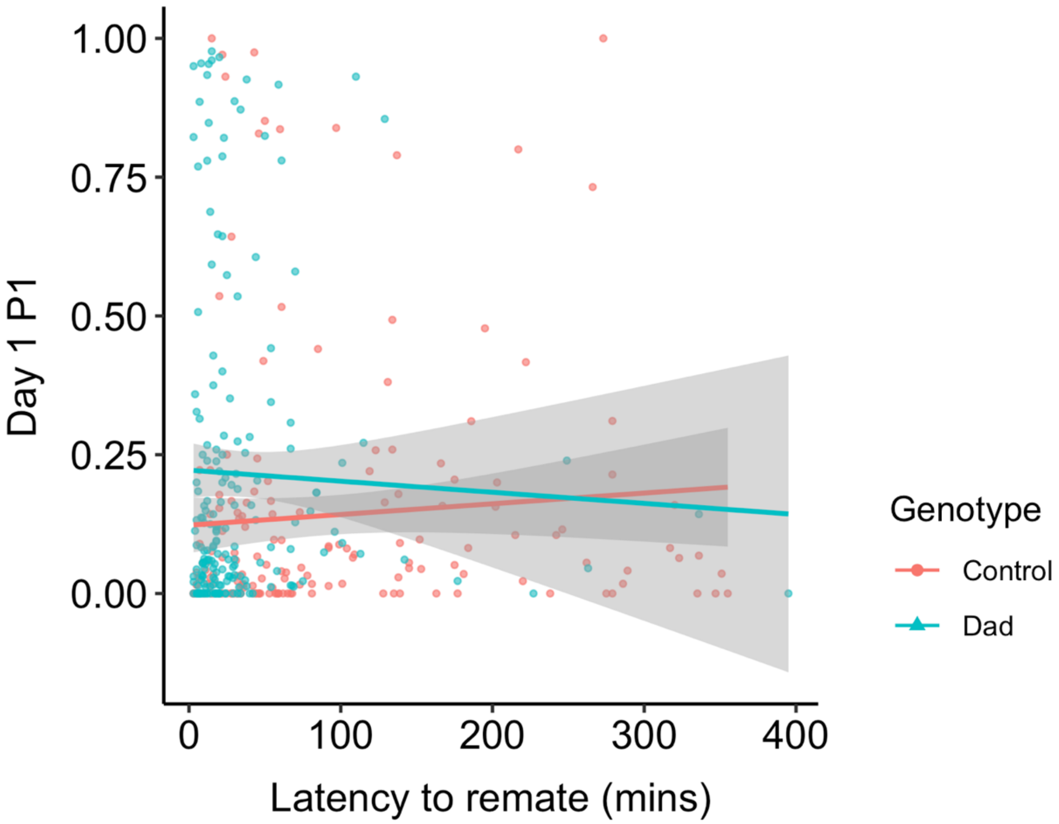


**Figure S3.** A regression of first male paternity share against latency to remating by Dad- (blue) and control- (red) mated females secondarily mated to a standardised competitor. $n_{Dad}$=190, $n_{control}$=173, pooled across 6 blocks. Confidence intervals drawn at the 95% level. Significant effect of genotype (*p*=0.001), but not of latency (*p*=0.608) nor the interaction between them (*p*=0.567).

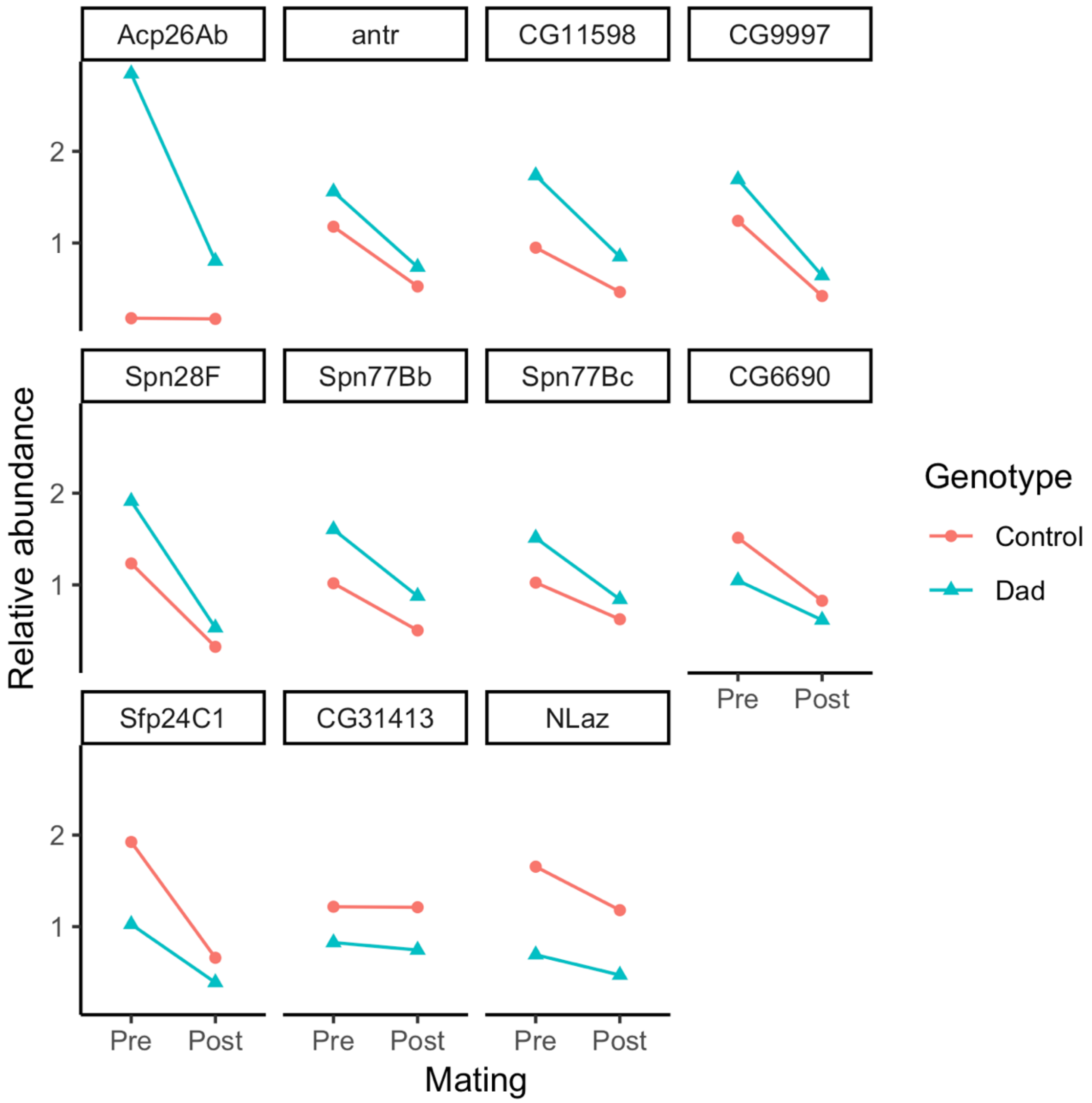


**Figure S4.** Abundance profiles of SFPs identified as differentially-abundant in relation to genotype (FDR $p<0.05$). Each point represents an average across the 5 replicates in relation to genotype and mating status. The abundance values are relativized by means-centring and averaging across replicates.

**Table S1.** Summary statistics from a PCA conducted on detected SFPs. (A) The variance, eigenvalue, and loadings associated with the first four principal components (PCs). (B) The output from linear models fitted to each of the first three PCs, using the measured variables of genotype (Dad or control), mating status (mated or pre-mating), and replicate (5 in total). Significant associations at the $p<0.05$ levels are given in red.

# A

| | *PC1* | *PC2* | *PC3* | *PC4* |
|---|---|---|---|---|
| *Variance explained (%)* | 60.76 | 7.77 | 6.15 | 4.35 |
| *Eigenvalue* | 53.47 | 6.84 | 5.41 | 3.83 |
| *No. positive loadings* | 82 | 42 | 47 | 44 |
| *No. negative loadings* | 6 | 46 | 41 | 44 |

# B

| *PC1* *Effect* | *Df* | *Sum of sq* | *RSS* | *F* | *P* |
|---|---|---|---|---|---|
| *Genotype*Mating* | 1 | 3.6962 | 11.3870 | 5.7672 | 0.0334 |
| *Genotype* | 1 | 18.56 | 29.95 | 21.190 | 0.0004 |
| *Mating* | 1 | 971.50 | 982.88 | 1109.1 | <0.0001 |
| *Replicate* | 4 | 14.52 | 25.90 | 4.1433 | 0.0222 |

| *PC2* *Effect* | *Df* | *Sum of sq* | *RSS* | *F* | *P* |
|---|---|---|---|---|---|
| *Genotype*Mating* | 1 | 1.1368 | 9.4912 | 1.6329 | 0.2255 |
| *Genotype* | 1 | 102.42 | 111.91 | 140.28 | <0.0001 |
| *Mating* | 1 | 1.8380 | 11.329 | 2.5176 | 0.1366 |
| *Replicate* | 4 | 16.200 | 25.691 | 5.5474 | 0.0079 |

| *PC3* *Effect* | *Df* | *Sum of sq* | *RSS* | *F* | *P* |
|---|---|---|---|---|---|
| *Genotype*Mating* | 1 | 2.0705 | 71.065 | 0.3601 | 0.5596 |
| *Genotype* | 1 | 4.3395 | 75.404 | 0.7938 | 0.3891 |
| *Mating* | 1 | 0.0131 | 71.078 | 0.0024 | 0.9616 |
| *Replicate* | 4 | 27.418 | 98.482 | 1.2539 | 0.3370 |

**Table S2.** SFPs detected as significantly differentially-abundant in response to genotype. q-values are calculated by applying a tail-based FDR correction to *p*-values obtained from a linear model iterated over each detected protein. q-values are given both for the effect of mating status (pre-mating/mated) and the genotype (Dad/control). Fold changes are given on a $\log_2$ scale and calculated for each genotype comparison within a mating status. Dad value is subtracted from the control. Therefore, positive values indicate greater abundance in controls. The transfer value is calculated by subtracting the Dad pre/post-mating fold change from the control pre/post-mating fold change. Therefore, positive values indicate greater transfer of an SFP to females in controls. Functional information associated with each protein's FlyBase entry is provided.

| | qval | | Log fold change | | | | |
|---|---|---|---|---|---|---|---|
| Protein | Genotype | Mating | Virgin | Mated | Transfer | Functional class | Predicted function |
| Acp26Ab | 0.0001 | 0.0252 | -3.9921 | -2.2305 | -1.7616 | Peptide/Prohormone | Post-mating behavior |
| antr | 0.0099 | 0.0001 | -0.4067 | -0.4717 | 0.0650 | CRISP | Defence response |
| CG11598 | 0.0001 | 0.0001 | -0.8491 | -0.8736 | 0.0245 | Acid lipase | Lipase activity |
| CG31413 | 0.0021 | 0.4695 | 0.5486 | 0.6916 | -0.1430 | Thioredoxin | Protein modification process |
| CG6690 | 0.0204 | 0.0001 | 0.5501 | 0.4407 | 0.1094 | Thioredoxin | Protein modification process |
| CG9997 | 0.0063 | 0.0001 | -0.4432 | -0.5971 | 0.1539 | Serine protease | Proteolysis |
| NLaz | 0.0001 | 0.0163 | 1.3765 | 1.3397 | 0.0368 | Lipocalin | Lipid metabolic process |
| Sfp24C1 | 0.0001 | 0.0001 | 0.9107 | 0.7763 | 0.1344 | Serine protease | Endopeptidase inhibitor |
| Spn28F | 0.0011 | 0.0001 | -0.6338 | -0.7392 | 0.1054 | Serpin | Negative regulation of proteolysis |
| Spn77Bb | 0.0003 | 0.0001 | -0.6584 | -0.8108 | 0.1524 | Serpin | Negative regulation of proteolysis |
| Spn77Bc | 0.0363 | 0.0002 | -0.5505 | -0.4633 | -0.0871 | Serpin | Negative regulation of proteolysis |

**Table S3.** Non-SFPs detected as significantly differentially-abundant in response to genotype. q-values are calculated by applying a tail-based FDR correction to *p*-values obtained from a linear model iterated over each detected protein. q-values are given both for the effect of mating status (pre-mating/mated) and the genotype (Dad/control). Fold changes are given on a $\log_2$ scale and calculated for each genotype comparison within a mating status. Dad value is subtracted from the control. Therefore, positive values indicate greater abundance in controls. Functional information associated with each protein's FlyBase entry is provided.

| | | qval | | Log fold change | | | |
|---|---|---|---|---|---|---|---|
| Protein | Synonym | Genotype | Mating | Virgin | Mated | Functional class | Predicted function |
| 34F4T | CG6084 | 0.0144 | 0.5982 | -0.5304 | -0.2470 | Aldo-keto reductase | Oxidoreductase activity |
| CG8628 | | 0.0234 | 0.5915 | -1.3997 | -0.9496 | Acyl-CoA binding | Fatty-acyl-CoA binding |
| Arc1 | CG12505 | 0.0145 | 0.3881 | 0.7358 | 0.6966 | ARC/ARG3.1 family | mRNA binding, vesicle transport |
| Ars2 | CG7843 | 0.0146 | 0.0402 | -0.5872 | -0.4427 | Protein binding | RNA processing |
| BP1025 | CG34002 | 0.0381 | 0.0001 | -0.2350 | -0.3303 | CRISP | |
| CCT3 | CG8977 | 0.0003 | 0.0001 | 0.5754 | 0.5030 | TCP-1 chaperonin | ATP-binding |
| CG12262 | Mcad | 0.0292 | 0.4068 | 0.1925 | 0.1962 | Acyl-CoA dehydrogenase | Acyl-CoA dehydrogenase activity |
| CG1532 | | 0.0185 | 0.0001 | -0.2595 | -0.2447 | | |
| CG34034 | BP1088 | 0.0006 | 0.0001 | 0.9092 | 0.7807 | | Ejacualtory duct protein |
| CG7408 | | 0.0212 | 0.2767 | 0.4288 | 0.3982 | Sulfatase | Sulfuric ester hydrolase activity |
| CG7966 | | 0.0002 | 0.6415 | -1.4620 | -1.5465 | | Selenium binding |
| CG8303 | | 0.0197 | 0.6473 | -0.3672 | -0.4004 | Acyl-CoA binding | Fatty-acyl-CoA binding |
| Chd64 | CG14996 | 0.0350 | 0.6093 | -0.3380 | -0.1718 | Actin binding | Muscle contraction; JH response element |
| coro | CG9446 | 0.0483 | 0.0022 | -0.2318 | -0.1948 | | Somatic muscle development |
| Cpr | CG11567 | 0.0189 | 0.0670 | 0.2174 | 0.2890 | NADPH cytochrome P450 reductase | Oxidoreductase activity |
| GH26 | CG6287 | 0.0128 | 0.1044 | -0.2145 | -0.1897 | Oxidoreductase | NAD binding |
| GLaz | CG4604 | 0.0339 | 0.0001 | -0.2112 | -0.8633 | Lipocalin | Fatty acid binding |
| Hsp23 | CG4463 | 0.0274 | 0.5398 | 2.1436 | 1.8039 | Small heat shock protein | Protein folding |
| Hsp27 | CG4466 | 0.0016 | 0.6320 | 10.3680 | 12.0168 | Small heat shock protein | Protein folding |
| Idh | CG7176 | 0.0466 | 0.4891 | 0.2491 | 0.2592 | Oxidoreductase | Isocitrate dehydrogenase activity |
| Lsp2 | CG6806 | 0.0237 | 0.5855 | -0.3754 | -0.5225 | Hemocyanin | Amino acid storage |
| Ncc69 | CG4357 | 0.0192 | 0.0065 | -0.4123 | -0.4141 | Chloride transporter | Transmembrane transporter activity |
| NTPase | CG3059 | 0.0127 | 0.0244 | -0.2493 | -0.2697 | Nucleoside phosphatase | Diphosphatase activity |
| PPO1 | CG42639 | 0.0004 | 0.0939 | -0.7474 | -0.7863 | Phenoloxidase | Immune/Melanization |
| sphinx2 | CG32382 | 0.0140 | 0.0008 | -0.7809 | -0.9257 | Serine protease homolog | Proteolysis |
| tweek | CG42555 | 0.0001 | 0.1373 | 2.5610 | 1.9503 | Fragile site associated protein | Vesicle endocytosis |
